# Bayesian models for event-related potentials time-series and electrode correlations estimation

**DOI:** 10.1101/2022.08.02.502520

**Authors:** Simon Busch-Moreno, Xiao Fu, Etienne B. Roesch

## Abstract

Recent developments on Bayesian inference and prediction for effects on event-related potentials (ERPs) stem from a wide variety of methods, including machine learning, multilevel models, and others. However, few of these approaches make use of clear estimates of the ERP voltage across time and across space (scalp). In the present article, via an iterative process, we propose Gaussian random walk (GRW) models that can estimate voltage uncertainty across time and also provide correlation matrices across electrodes. We apply these models to real ERP data from a P3b paradigm as an example. We discuss results in terms of past and current literature of both ERP estimation and electroencephalography analysis in general.

## Introduction

Event-related potentials (ERPs) are generally understood as time-series (Cohen, 2014; Luck, 2014). However, the most commonly used analysis technique summarises a specified time-window in the form of average amplitudes to be analysed through a regression across groups and conditions, or similar analysis techniques (e.g. Luck, 2014). Alternatively, ERPs are analysed in the time domain, but as a series of t-tests, F-tests, or similar, performed at each sample point (time-point) of the baseline and epoch (e.g. permutation-based cluster tests; see Sassenhagen et al., 2019). The problem of the former approach is that the time-series nature of the ERP signal is lost when a window of time is summarised in a single statistic, therefore the analyses are not addressing a time-series problem anymore and the time-domain information is lost. The second approach attempts to extract information from the time-domain, but it suffers from numerous other problems (see Sassenhagen et al. 2019), such as requiring corrections for multiple comparisons, which is an ad-hoc statistical device (see Jaynes, 2003; Gelman, 2006; Gelman et al., 2014). Such ad-hoc devices are generally not satisfactory solutions, because they rarely resolve the frequentist problem of inflated type-I error due to large scale observations (see Efron, 2010). As an alternative, Bayesian approaches for detecting single trial ERPs are considered to be generally reliable, but current proposals operate on variational maximum likelihood (ML) or maximum a posteriori methods (MAP) (see Sanei, 2013; Sanei & Chambers, 2021). ML and MAP methods may be problematic, as they essentially provide just point estimates (i.e. they are not fully Bayesian, see Murphy, 2012). Mean-field based variational inference, may also be problematic for present purposes, as it is not particularly good at estimating correlations in the posterior (see van de Schoot et al., 2021), such as correlations between electrodes. Therefore, in the present article we attempt to provide a simple proof of concept, where ERPs can be estimated as time series. Our approach focuses on providing reliable uncertainty estimates on the ERP, as well as providing correlations between electrodes across the scalp.

To this aim, we implement a Bayesian model using multivariate Gaussian random walk (MGRW) prior on ERPs from all electrodes of an electroencephalography (EEG) recording. The idea behind using a MGRW is that EEG signals’ noise/error can be modelled via a Gaussian distribution (Cohen, 2014; Luck, 2014; Sanei, 2013); more precisely, a multivariate Gaussian (i.e. across all channels/electrodes) of a semi or quasi-stationary signal (Sanei, 2013; Sanei & Chambers, 2021). Gaussian distributions to represent the distribution of voltage’s noise are also assumed to be appropriate models for the synchronised firing of neural populations, such as in the post-synaptic potentials that produce ERPs (Sanei, 2013). Although a Gaussian process may be a more accurate model of the time series progression across the baseline-epoch, a MGRW provides a reasonable and sensible alternative, which also allows us to include a prior for the covariance matrix over electrodes, thus providing relevant information of scalp distributions for a model that is relatively easy to sample. The rationale is that a MGRW being a cumulative sum of Gaussians across the time-samples can provide good uncertainty of the ERP signal (voltage) across baseline and epoch at each electrode simultaneously.

In the present article, we describe three models: the first model consists simply of a multivariate Gaussian distribution for modelling the voltage across time-samples; the second is a MGRW model using an identity matrix as prior for covariance; and the third model consist of a MGRW model using a Lewandowski-Kurowicka-Joe (LKJ) prior for the covariance matrix. Models were implemented on data from an oddball task, where participants were instructed to listen to four Chinese Mandarin tones (Tone 1, Tone 2, Tone 3 and Tone 4), where Tone 4 was the target (25% of trials) and the other tones where non-targets. EEG data was collected from 32 electrodes and pre-processed with an automated pre-processing pipeline (details below), including independent component analysis (ICA) artefact removal, filtering and baseline correction. Full details about data collection and pre-processing of these EEG data are provided elsewhere (Fu, 2022). ERPs were obtained as the averages over trials and participants, which although not ideal allowed us to scale matrices down for improved sampling. Another drawback of our approach is that from a Bayesian perspective, pre-processing of signals is not necessary, because statistical error/noise can also be modelled and sampled from the posterior (see Jaynes, 2003). Having mentioned this, our aim is to provide a proof of concept for an analysis of ERP as time-series that can be used complimentarily to other Bayesian regression approaches (e.g. over average amplitudes from pre-specified timewindows), and which can be efficiently sampled with Hamiltonian Monte Carlo (HMC) methods using data from a conventional ERP analysis approach. Future implementation of more comprehensive models that do not discard information from the EEG signals may be required in specific contexts, such as models that can account for trial-by-trial time-series, or even models that directly detect end estimate events from raw signals. We believe the approach we describe provides a reasonable and sensible example of modelling ERPs as time-series.

## Models

Our first model uses an Identity matrix as covariance for the MGRW, simplifying the sampling process. But this also implies that we cannot obtain covariance/correlation matrices across electrodes.

### Model 1 Multivariate Gaussian Random Walk with covariance identity matrix

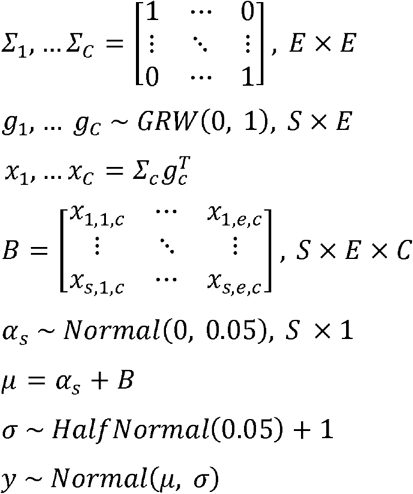

Where *S, E* and *C* are the same as in the previous model, *∑_c_* are four identity matrices (one per tone), *g_c_* are four GRWs priors for samples across electrodes, *x_c_* are four MGRWs (using the same reparameterization technique as previous model), *B* is a design matrix containing of all MGRWs across time, electrode and condition (tone), *α_s_* is a scalar of Gaussian distributions across time to provide stationary noise (as an intercept), and *μ* and *σ* are the parameters for a Gaussian distribution. Priors for *σ* and *α_s_* were determined via prior predictive checks. Note that we obtain each MGRW prior x by multiplying each GRW prior *g* with the corresponding matrix *Σ*, namely a form of non-centred reparameterization (see Betancourt & Girolami, 2015; McElreath, 2020).

It was not feasible to integrate groups in the present model, so we sampled two separate models for learners and non-learners (group 1 and group 2). Models were sampled using no U-turn sampling (NUTS), a form of Hamiltonian Monte Carlo (HMC) sampling, as implemented in PyMC (Salvatier et al., 2016). We used 4 chains with 1000 tuning steps and 1000 samples each, they had very good convergence with all Bayesian Fraction of Missing Information (BFMI) > 0.9, Effective Sample Size (ESS) > 2000, and 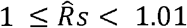. Our repositories contain detailed summaries of estimates (see data availability statement below). Model 1 provides sensible estimates of the signal. Figures 1 and 2 provide examples of posteriors from electrode Pz, from the learners and nonlearners groups respectively.

**Figure 1.**
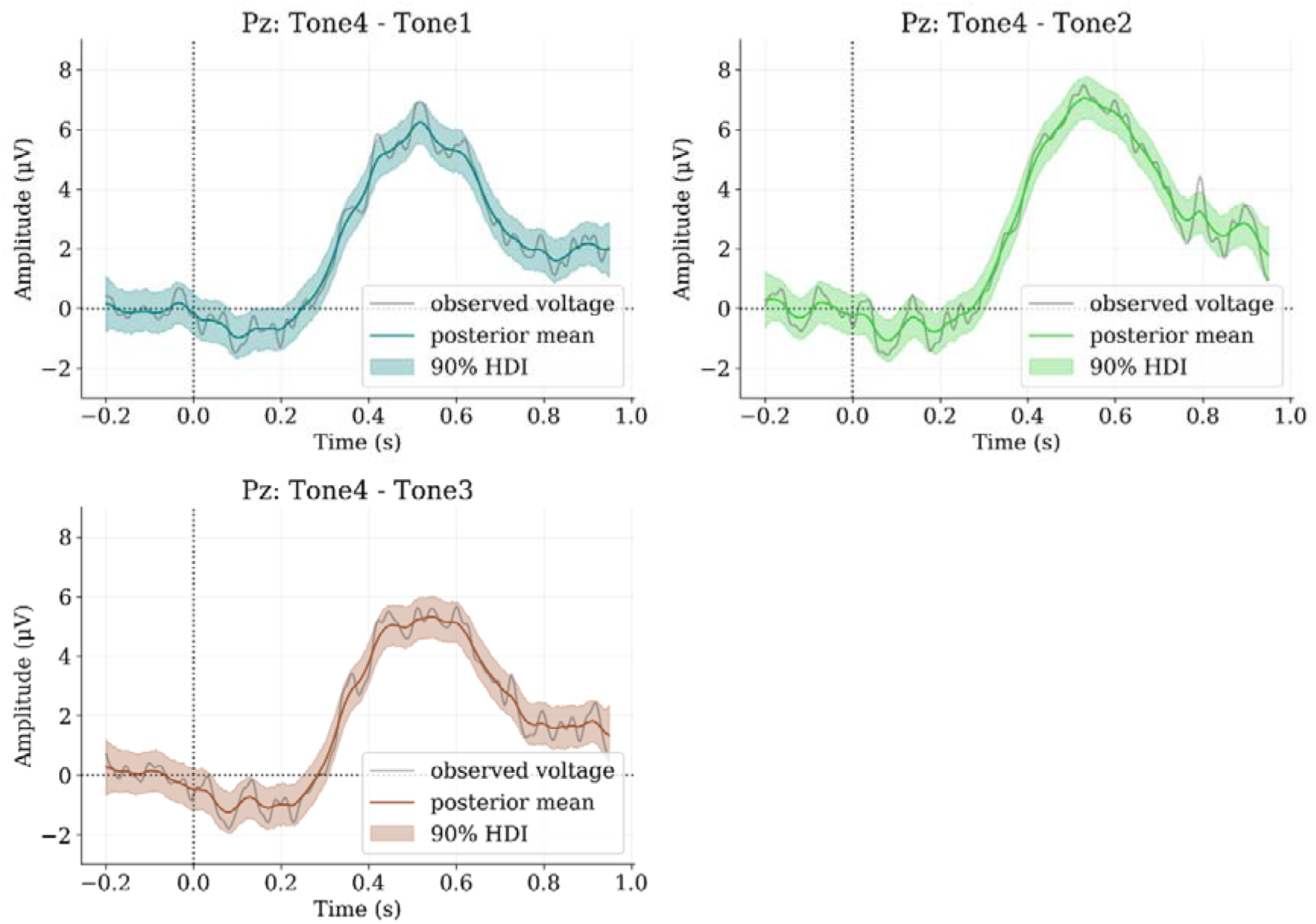
Model 1. Learners. Posterior difference waves between target tone and non-target tones. HDI: highest density interval.

**Figure 2.**
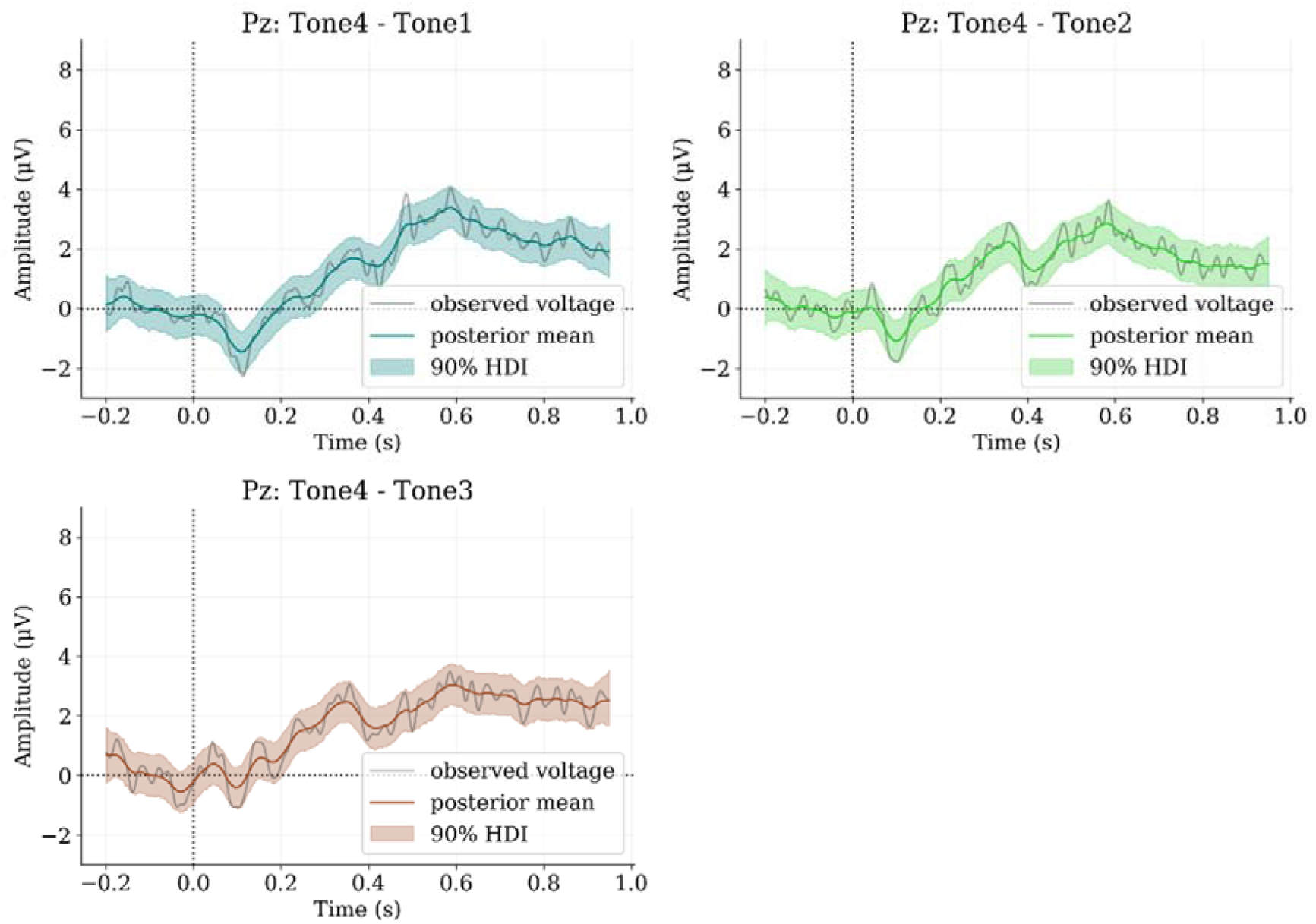
Model 1. Non-learners. Posterior difference waves between target tone and non-target tones. HDI: highest density interval.

Model 1 provides good estimates, but fails to provide estimates for electrodes correlations, which can be very relevant for a better topographical assessment of the ERPs. It is possible, however, to reparametrize a GRW as the product of standard Gaussian (*w*), a standard deviation parameter (*σ*) and the square root of sampled times (*t*), i.e. [0…*S*]/*f*, where *S* = 295 samples, and *f* = 256*hz* sampling frequency.

We thus implemented a second model, which has the advantage that it can estimate all groups and conditions at once, namely from a matrix of multivariate Gaussians as the expectation parameter of a Gaussian distribution.

### Model 2 vanilla Multivariate Gaussian

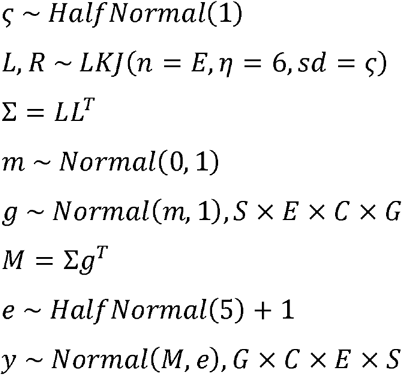

Where *S* = 295 *samples* (time-points), *E* = 32 *electrodes, C* = 4 *conditions* (tones), G = 2 *groups* (learners and non-learners). *L* corresponds to the Cholesky factor as derived from a Lewandowski-Kurowicka-Joe (LKJ) prior distribution (Lewandowski et al., 2009) which is also used to derive the electrodes correlation matrix *R*. LKJ distributions are usually the default for covariance/correlation matrices priors (see McElreath, 2020), as they generally perform better than distributions such as Wishart, which usually require adjustments to achieve equivalent performance (Wang et al., 2018). The covariance matrix Σ is derived via a Cholesky decomposition (*LL^T^*) and used in a non-centred reparameterization (∑*g^T^*) for the multivariate Gaussian prior *M*, which together with *e* are expectation and error parameters respectively of a Gaussian likelihood/sampling distribution *y*.

We sampled this model as Model 1. The model sampled relatively well with all BFMI > 0.8, all ESS > 700, and 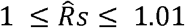. Our repositories contain detailed summaries of estimates (see data availability statement below). Although the model provides reasonable estimates of the signal, there may be a tendency to overfit and underestimate voltages. Figures 3 and 4 below shows examples of posteriors from the maximum amplitude electrode Pz from the learners group. Figure 5 shows 32 posterior means correlations () respect to Pz, from the LKJ correlation matrix over all conditions and groups.

**Figure 3.**
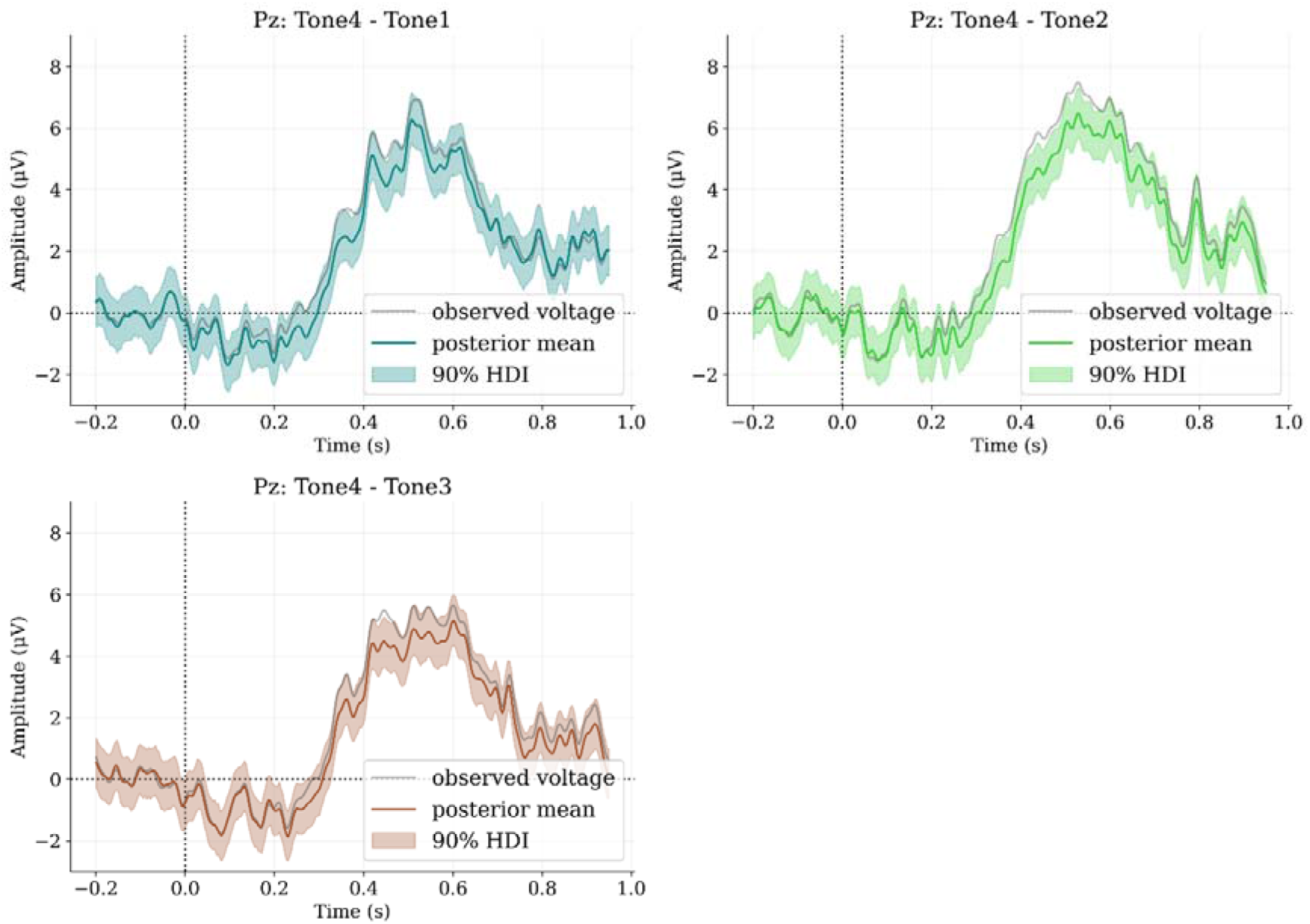
Model 2. Learners. Posterior difference waves between target tone and non-target tones. HDI: highest density interval.

**Figure 4.**
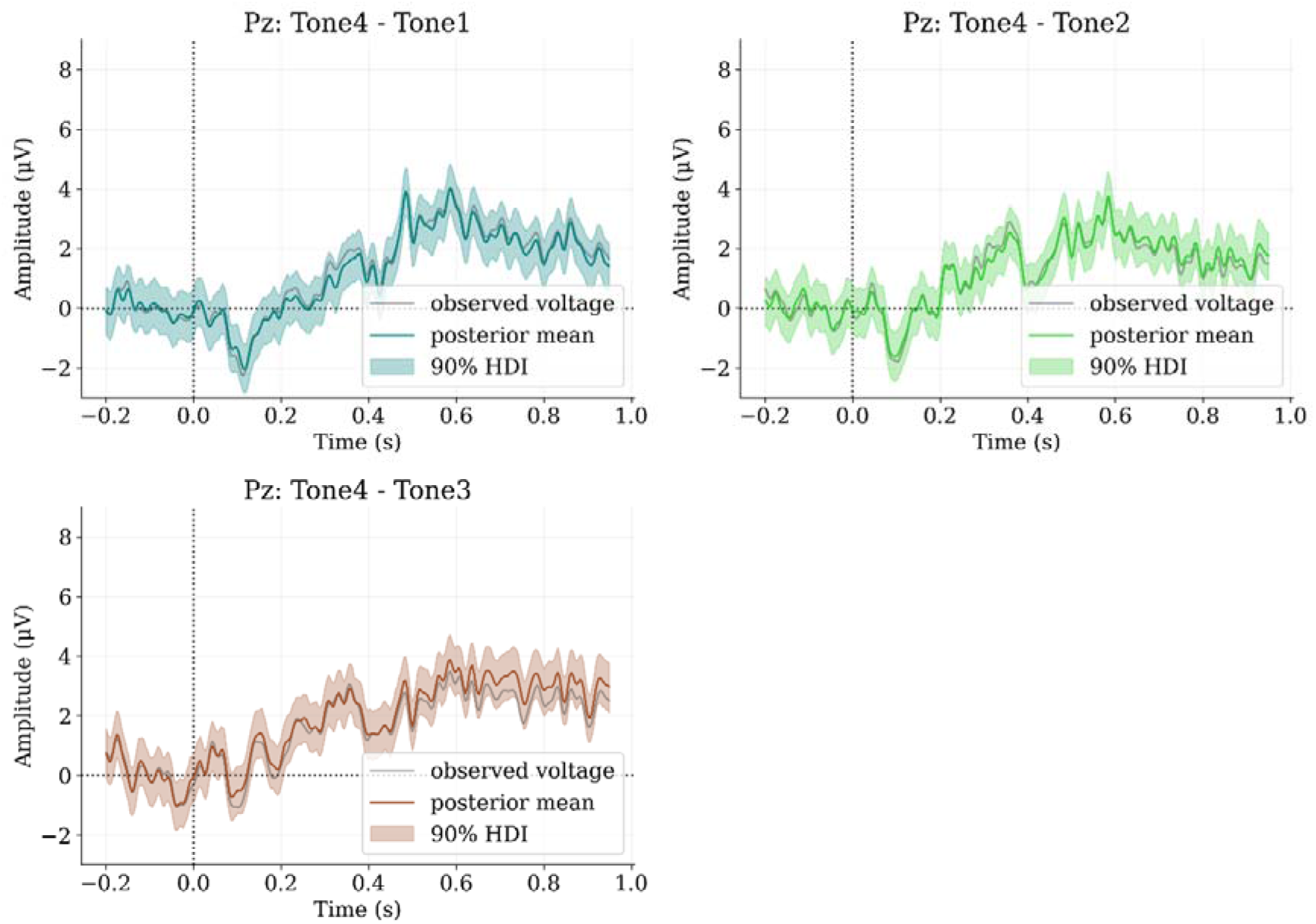
Model 2. Non-learners. Posterior difference waves between target tone and non-target tones. HDI: highest density interval.

**Figure 5.**
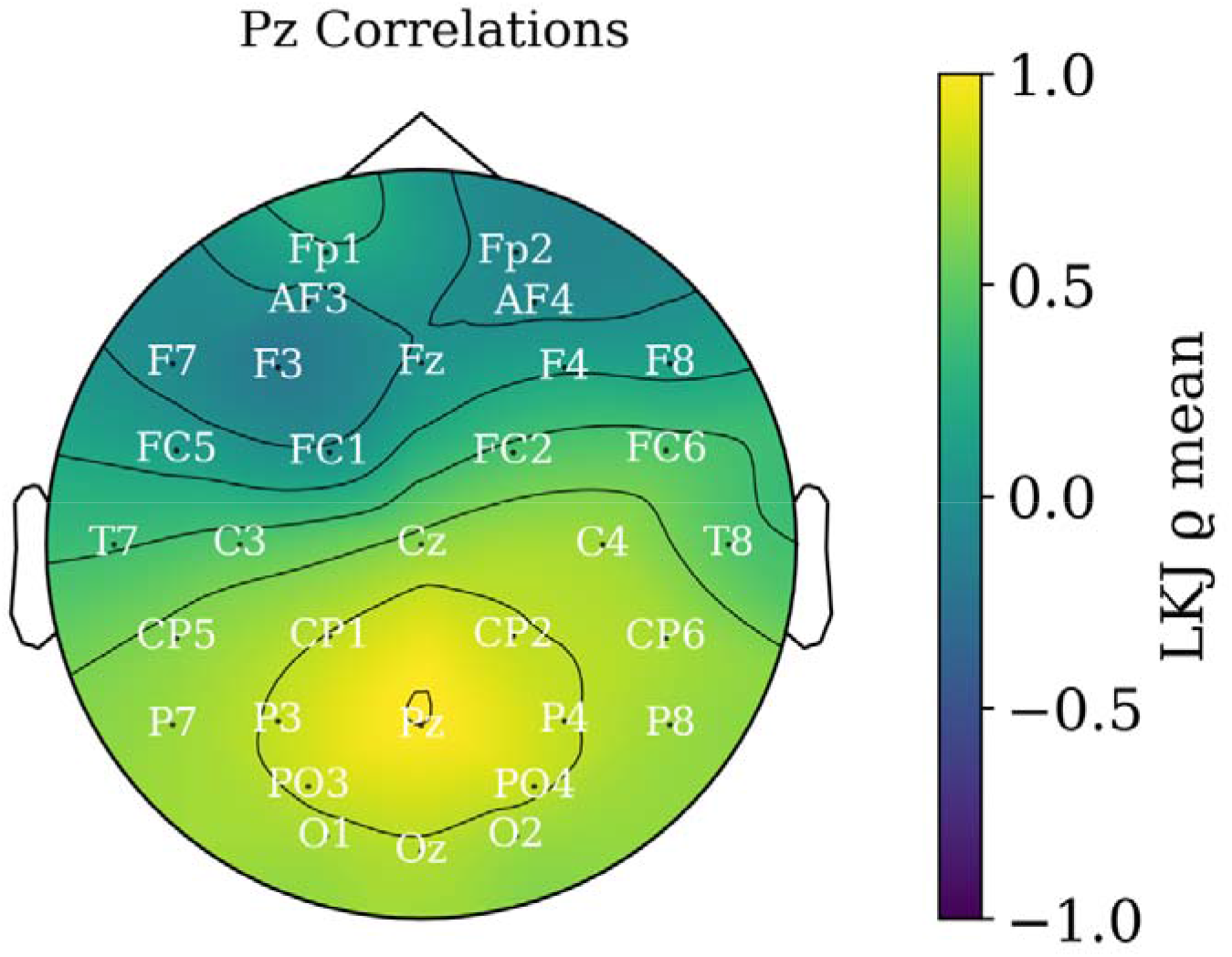
Model 2. Posterior mean correlations from LKJ prior on multivariate Gaussian distribution over time-samples, electrodes, conditions and groups.

Model 2 provides reasonable uncertainty estimates from groups and conditions, and correlations across electrodes confirm a cluster of associated electrical activity on the scalp (i.e. electrodes highly or lowly correlated to Pz, the maximum P3b voltage). However, voltages may not be independent across time-samples, as assumed by the model, which may be contributing to less accurate estimates and uncertainty, and may require a more sophisticated autoregressive model.

Thus, we attempt to enhance Model 2 by re-implementing a MGRW, but expressed as the product between a standard Gaussian distribution, a standard deviation parameter and the square root of sampled times. Which has a well-known proof: If 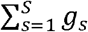 is a cumulative sum, such that: *g_s_, g_s_* + *g*_*s*+1_, *g_i_* + *g*_*s*+1_ + *g*_*s*+2_..., and all *g_s_* are distributed as a Gaussian distributions with mean 0 and variance *σ*^2^, and they are independent; then, by the properties of Gaussian distributions (see Lemons, 2002): *g_s_* + *g*_*s*+1_ = *N*(0, *σ*^2^) + *N*(0, *σ*^2^) = *N*(0 + 0, *σ*^2^ + *σ*^2^) = *N*(0, 2*σ*^2^) if all *σ*^2^ are equal. Therefore, by induction, 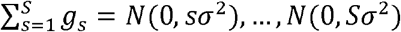, with *s* = 1,..., *S*. Therefore, we can parametrise via the standard deviation: 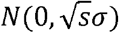 and reparametrize to a non-centred parametrization (see Betancourt & Girolami, 2015; McElreath, 2020): 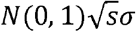, assuming zero location. So, if *w_s_* are standard independent Gaussian distributions such that *w_s_* ~ *N*(0, 1), our Gaussian random walk prior would be 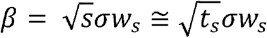 (see also: Caflisch, 2003). Adding one more dimension to *w_s_* we have 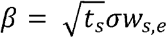, with *e* = 1,..., *E*. Then we can parametrise the MGRW as ∑*β*, where Σ is covariance matrix of shape *E* × *E* (as in Model 1). With this in mind, Model 3 reads as follows:

### Model 3 Multivariate Gaussian Random Walk with LKJ prior

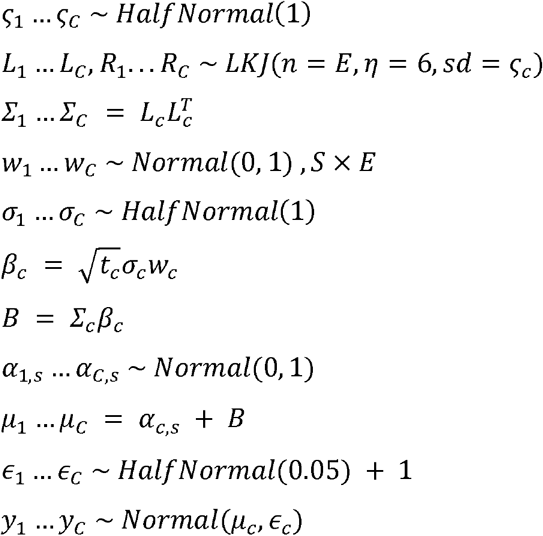

Where S = 295, E = 32, and C = 4; *L_c_* and *R_c_* are four Cholesky factors and correlation matrices as derived from the LKJ prior, *Σ*_1_ are four covariance matrices determined via a Cholesky decomposition, *w_c_* are four standard Gaussian distributions priors of shape S by E, *σ_c_* are four half-normal priors for standard deviations, *t_c_* are the sampled times (not to be confused with time-samples), *β_c_* are four GRWs priors for ERPs at all electrodes, *α_c,s_* are four priors for intercepts (stationary Gaussian noise) across samples, and *μ_c_* and *ϵ_c_* are priors for the estimate and standard deviation of Gaussian sampling distributions *y_c_*.

Model 3 was sampled as Model 1, but convergence was not ideal. All BFMI > 0.9, but although most ESS > 1000 16 parameters presented ESS below 1000 (though over 200), but 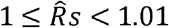. (Our repositories contain detailed summaries of estimates: see data availability statement below). These minor convergence issues may still imply that other reparameterization techniques may be required or maybe even the redefinition of some priors. Even so, the model provides reasonable estimates, as shown by Figures 6 and 7 portraying posteriors from Pz from learners and non-learners models respectively.

**Figure 6.**
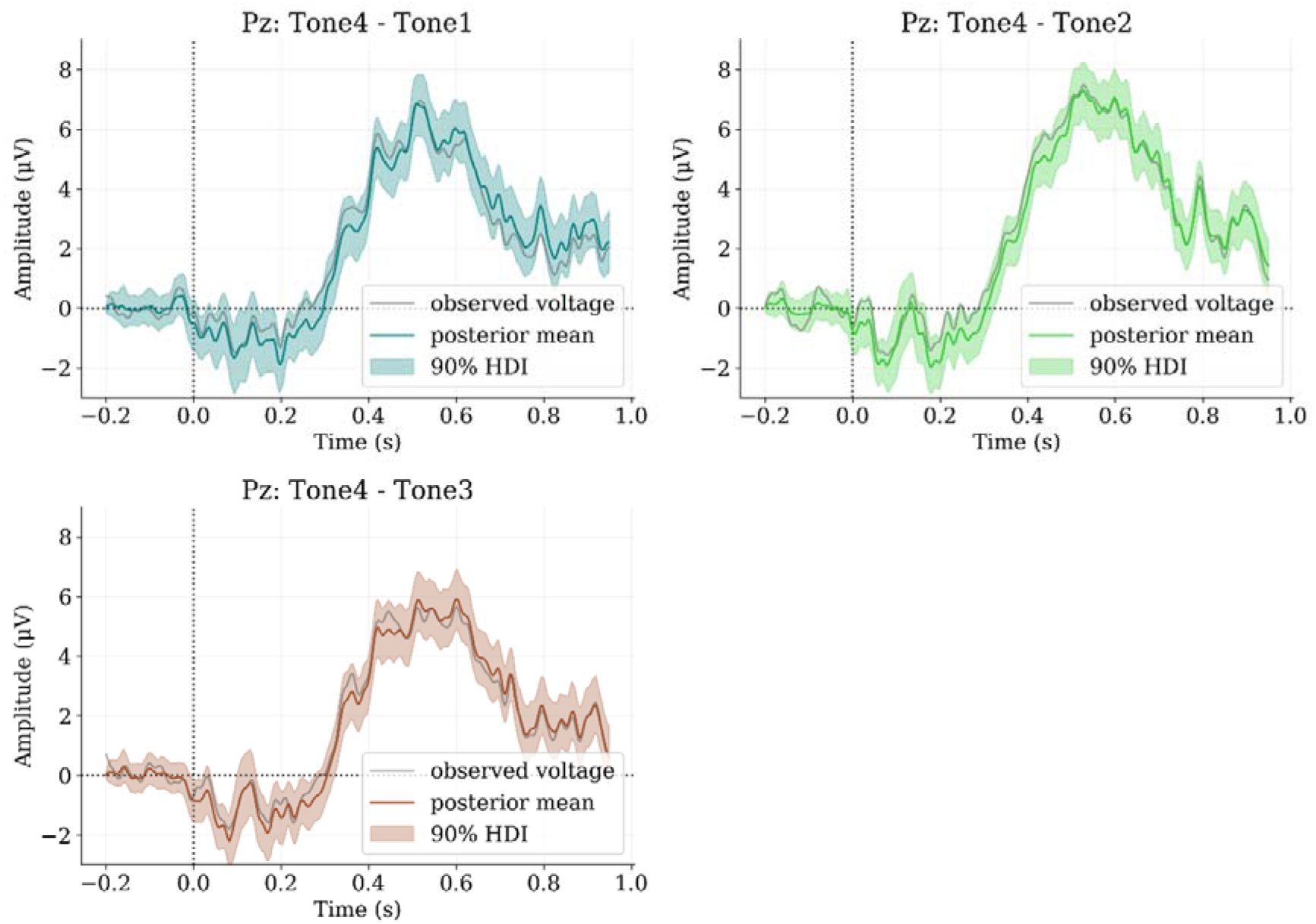
Model 3. Learners. Posterior difference waves between target tone and non-target tones. HDI: highest density interval.

**Figure 7.**
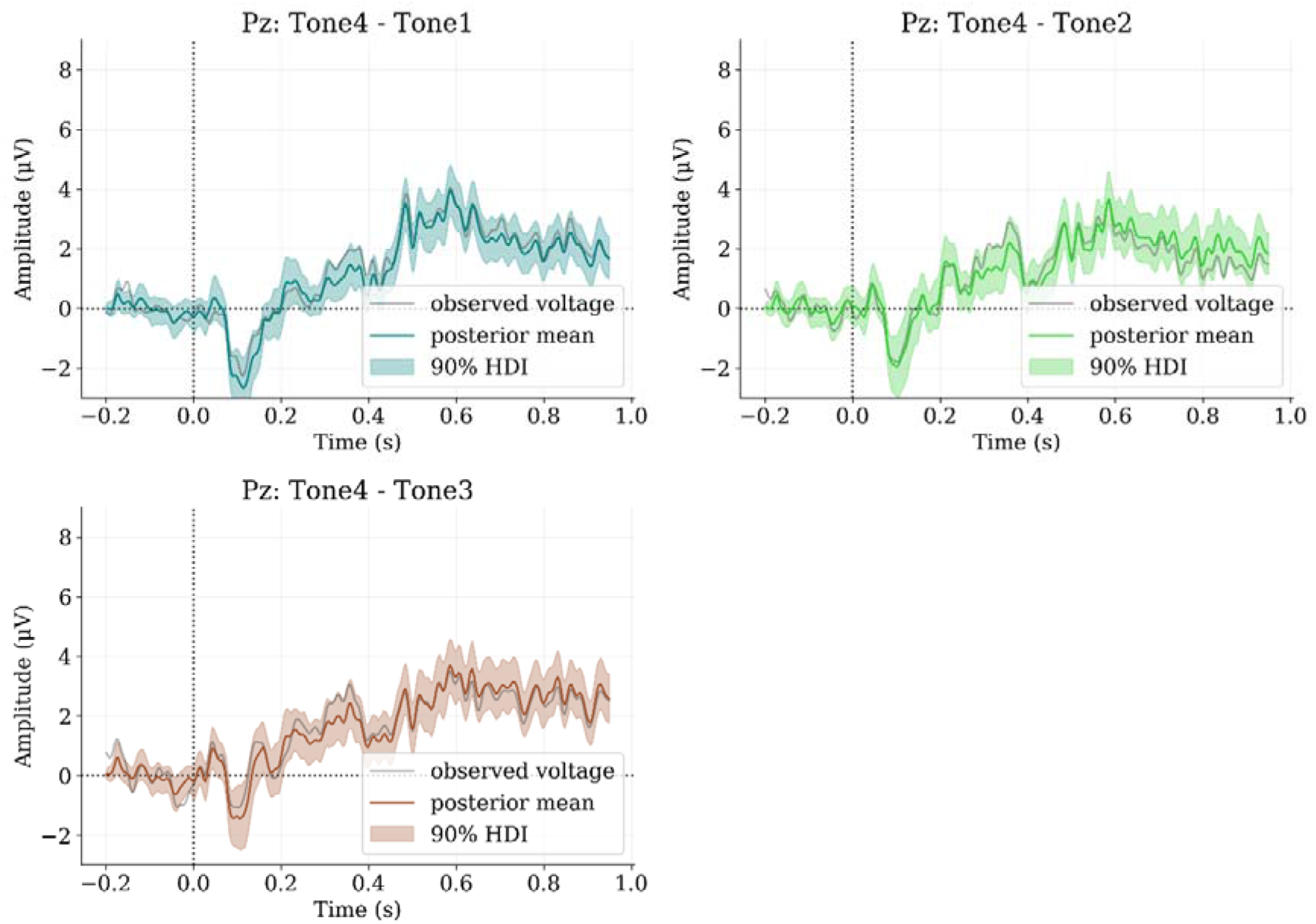
Model 3. Non-learners. Posterior difference waves between target tone and non-target tones. HDI: highest density interval.

Importantly, Model 3 can provide estimates of correlations between electrodes via an LKJ prior for each condition. Figures 8 and 9 show posterior means from Pz from each condition (tone) from learners and non-learners groups respectively. Even though estimates are better than Model 1, they do not seem as good as Model 2. However, correlations provide information that seems reliable, where a cluster of electrodes is highly associated to Pz (maximum P3b amplitude) for the learners group at the Tone 4 condition only, which is consistent with the expected P3b voltage variation.

**Figure 8.**
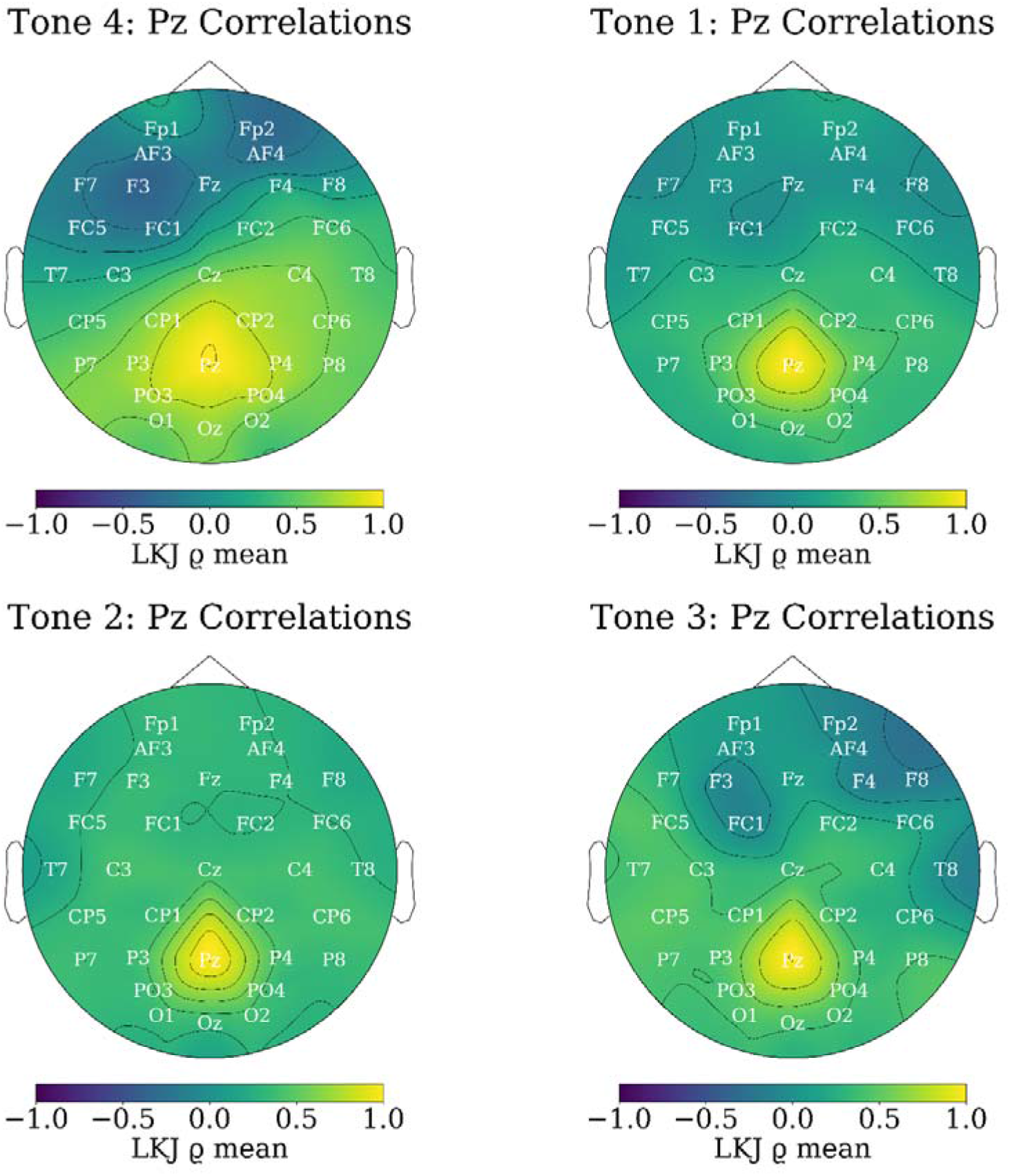
Model 3. Learners. Posterior mean correlations from LKJ prior on multivariate Gaussian random walk (MGRW) over time-samples, electrodes, and conditions.

**Figure 9.**
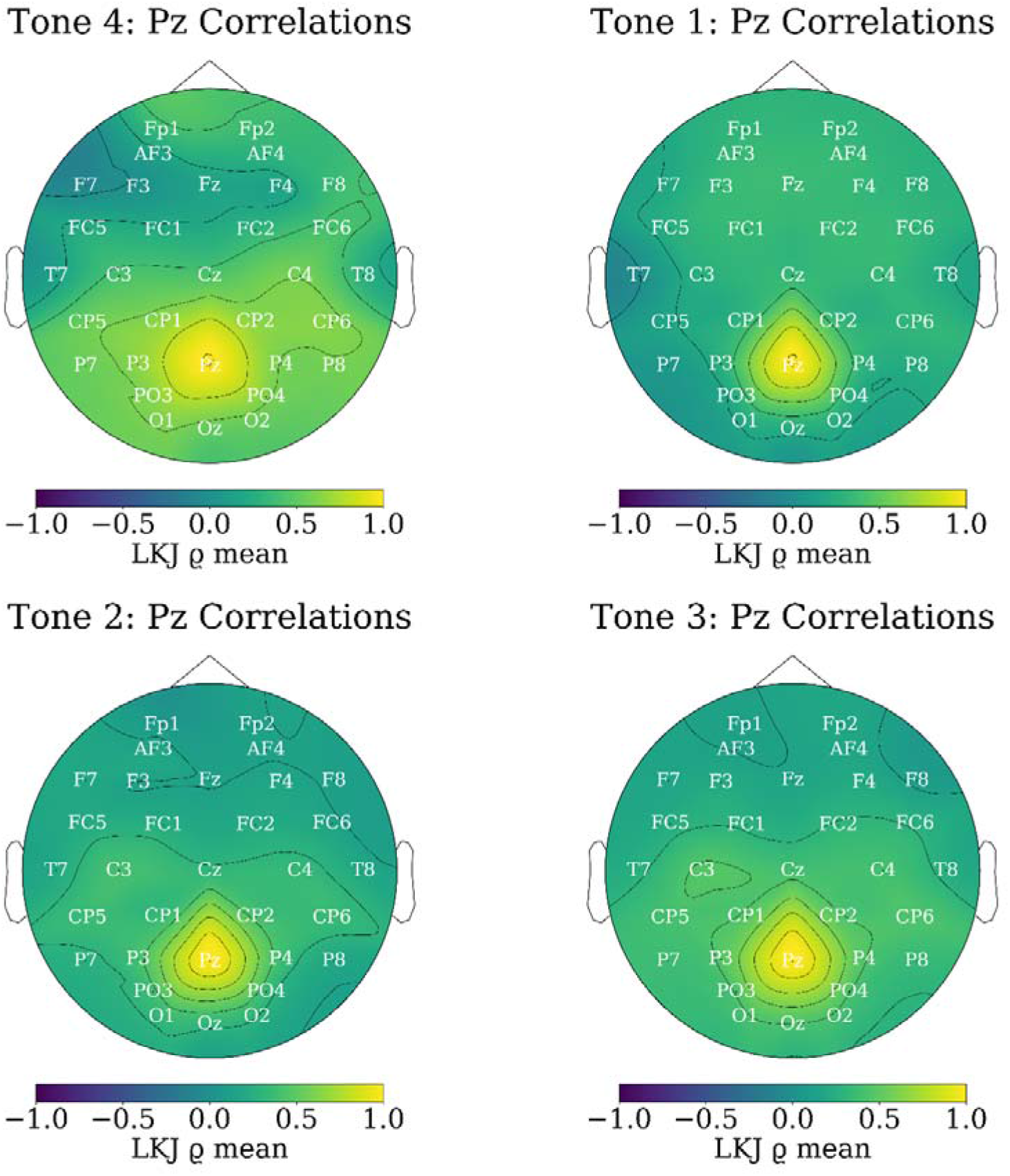
Model 3. Non-learners. Posterior mean correlations from LKJ prior on multivariate Gaussian random walk (MGRW) over time-samples, electrodes, and conditions.

## Discussion

Altogether, present models show that ERPs can be sensibly modelled as time series via variations on Gaussian priors, from simple models assuming a stationary signal (Model *2*) to more nuanced models using Gaussian random walks (GRW) to model non-stationary elements of the signal (Model 1 and Model 3). In addition, our models demonstrate that it is possible to capture electrodes covariances and correlations, which prove to be not only relevant but crucial to understand ERP distributions on the scalp and how they can be reflected by the associations between electrodes across the time-course of baseline and epoch. Even so, the main weakness of present models is that they require voltages averaged over trials and participants. This makes the modelling of errors/noise even more feasible due to the central limit theorem, but this averaging procedure also losses information from participants and trials variation.

Nevertheless, it is clear from present models that the assumptions of Gaussian-distributed noise (Sanei, 2013) across the time-course of ERP voltages is reasonable. Despite the lack of information from trials and/or participants, models provide relevant information on the uncertainty of the estimated signal. Predictions from the posterior, however, indicate that posterior predictive distributions have high uncertainty. Even though our models are not strongly intended to be predictive instruments, the future development of more efficient generative models is important. There are examples of these models in past literature (Sanei, 2013), which may be harder to sample from a Hamiltonian Monte Carlo (HMC) perspective. Thus, the exploration of these more intricate models may also require the use of variational inference which is also able to acknowledge correlations within the posterior, or similar methods (e.g. Semenova et al., 2022).

In the meantime, we demonstrate models of the sort can be useful tools to leverage the unfolding nature of ERPs as time-series of voltages in a feasible way, especially if used complimentarily with multilevel regressions over voltages averaged over time-windows of interest (e.g. Busch-Moreno et al., 2021; Fu, 2022). These techniques lose temporal information, but they retain the information from trials and participants. Also, if the appropriate distributions for priors and sampling are used, these models can be quite unsensitive to outliers, providing a good way for “cross-examining” present models. It is important to emphasise, however, that simpler models are not necessarily better, even if they prove more feasible for sampling. This is why we avoid misguided appeals to Ockham’s razor principles, as it is usually confused in its directions, where simpler models do not necessarily imply better explanations, but better explanations often imply simpler model (Jaynes, 2003). We emphasise “simpler” rather than simple, as simplicity is not an absolute category but is always respect to something. In this sense, present models may be too simple, at the cost of relevant information.

Therefore, the next step is to construct models and parametrisations procedures to cope with enough levels of complexity to do justice to voltage variation across time, electrodes, participants and trials. Previous attempts have focused on tackling combinations of these variables, but on all of them simultaneously and often blindly. For instance, ICA has been used for detecting single trial ERPs (see Sanei, 2013 pp 409-419), providing only a coarse perspective. Other techniques attempt to approach more than one parameter, such as participants, trials and electrodes (e.g. Busch-Moreno et al., 2021; Nicenboim et al., 2020), but miss the unfolding dimension of time and the correlation matrices between relevant parameters (i.e. electrodes or conditions). Acknowledging that loss of information was the focus of the present approach. Although this provides good results, it also makes evident that further development of models is required. These must also overcome some issues with past Bayesian models (e.g. Sanei, 2013 pp. 443-455) that rely on techniques such as MAP, which are no reliable for sampling complex posterior distributions, especially when implying correlations. Recent developments on Bayesian ERP modelling, including Bayesian Machine Learning for EEG analysis (Wu et al., 2016), indicate that this progress is feasible and may be achieved soon.

In conclusion, we presented three models that take advantage of the Gaussian noise assumption on ERP modelling. Models perform generally well, and despite their limitations provide a good way for estimating ERP voltages across the time-line (baseline and epoch). Also, two of these models provide estimates for correlations between electrodes, and show how relevant these are for a proper understanding of the ERP distributions. We discuss possible lines for future development of ERP models, where we emphasise the importance of modelling all relevant aspects of the signal, including time-course, electrodes, participants, conditions and trials. This may also lead to the development of modelling techniques of the raw EEG signal, altogether avoiding the conventional pre-processing of ERP signals that can be quite aggressive in its use of filters, removal of components via ICA, and baseline correction. Thus, we see present models as a relevant starting point which we can use to boost future, more nuanced, modelling of ERP signals.

## Supporting information

Annex 1

Annex 2

## Authors Contributions

Contributions to the study were as follows. **Simon Busch-Moreno:** conceptualisation, design, software, statistical methodology, analysis, visualisation, writing. **Etienne Roesch:** administration, design, writing, revision. **Xiao Fu:** experimental methodology, data collection, pre-processing, revision.

## Data Statement

All data is publicly available at: https://osf.io/h7kvm/. All scripts are available in GitHub. Model 1: https://github.com/ebrlab/Bayesian_MGRW_ERP. Model 2: https://github.com/ebrlab/BayesianERPsimple. Model 3: https://github.com/ebrlab/Bayesian_MGRW_LKJ_ERP.

## Ethics Statement

EEG data was collected from human participants as part of Xiao Fu’s dissertation project with ethical procedures according to UCL’s regulations for project: SHaPS-2018-BE-026. In accordance with Helsinki declaration for human research and GDPR.

## Acknowledgements

We thank Harrison Curtis, OHBM conference participants who visited our poster, and members of PyMC Discourse who answered our questions, for the fruitful discussions.

