## Supplementary material for "Bayesian models for event-related potentials time-series and electrode correlations estimation": Annex 1

Present supplementary materials provide additional results, showcasing posterior distributions as topographies. ERP waves from all electrodes can be found in our OSF and GitHub repositories (see data availability statement).

*Model 1 Posterior Topomaps*

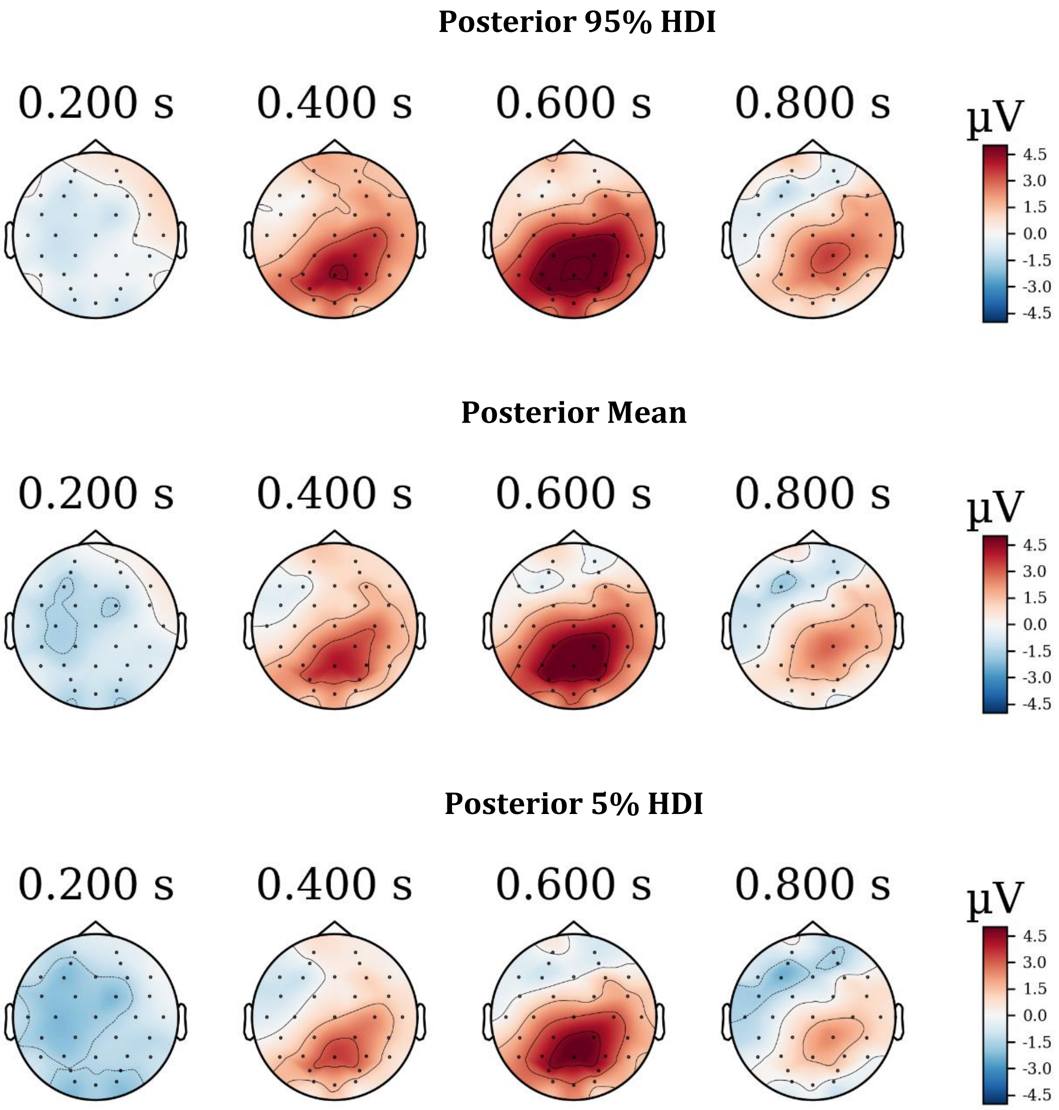


***Figure 1.*** Learner’s posterior distributions. Plots show estimated amplitude across the scalp (32 electrodes). HDI: highest density intervals

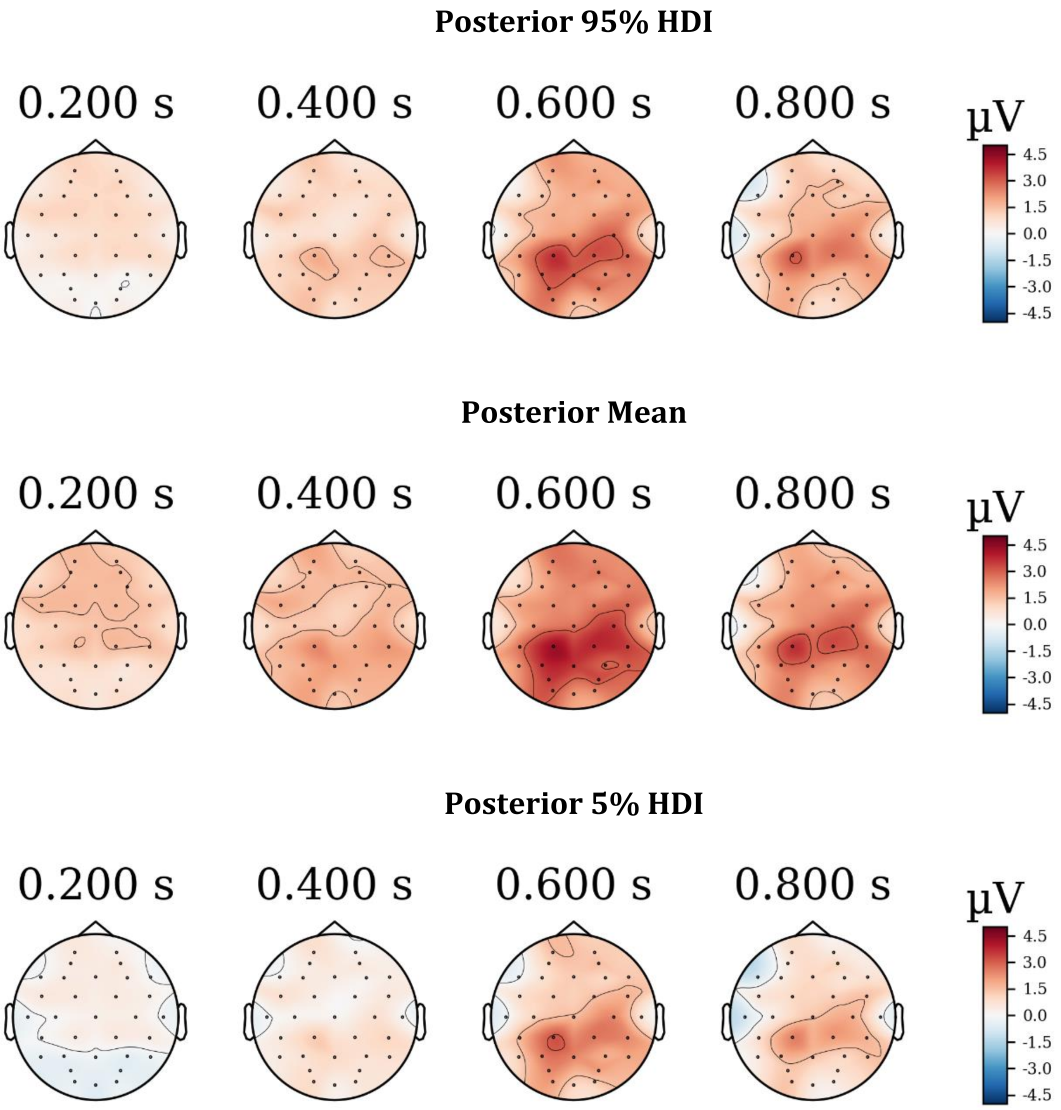


***Figure 2.*** Non-Learner's posterior distributions. Plots show estimated amplitude across the scalp (32 electrodes). HDI: highest density intervals

*Model 2 Posterior Topomaps*

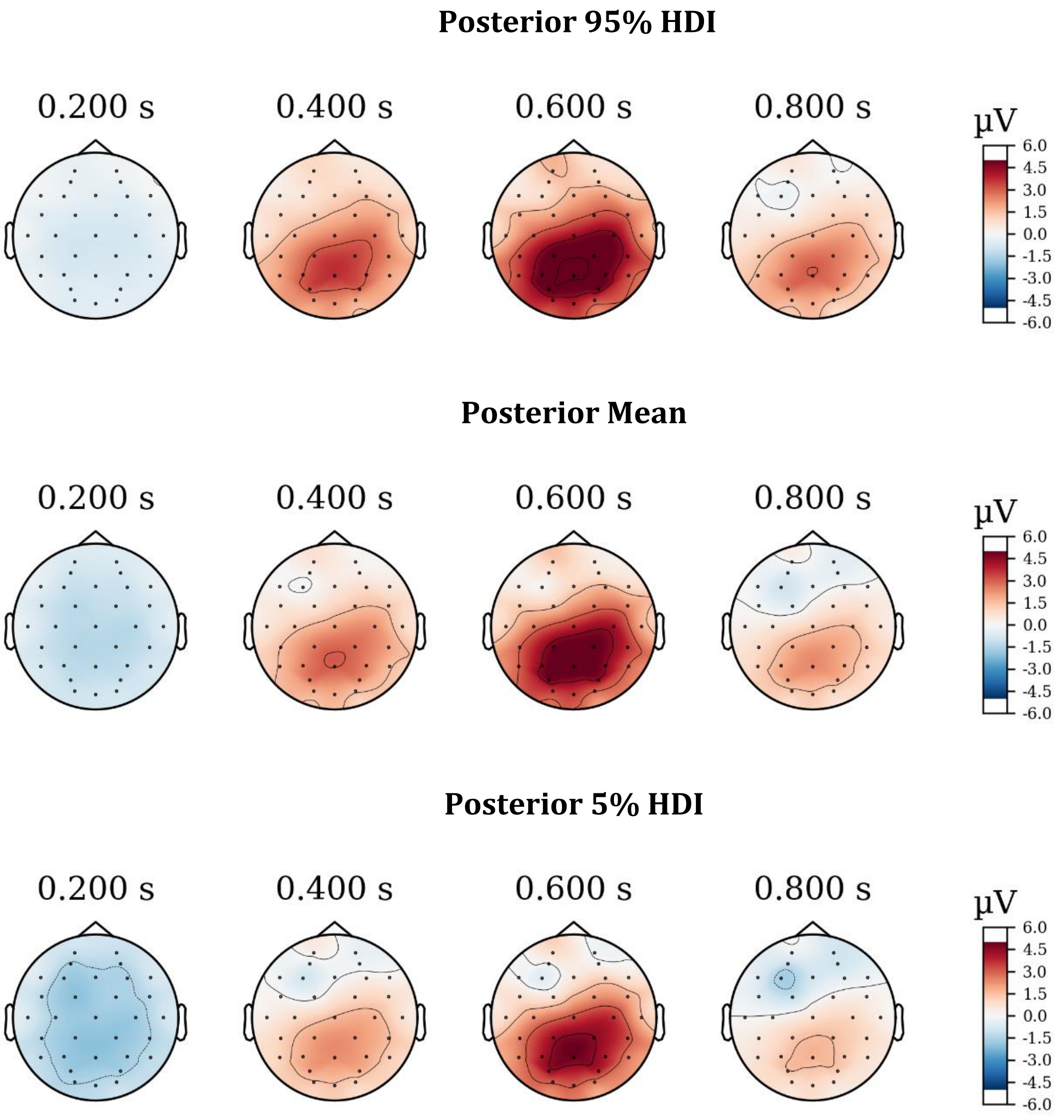


***Figure 3.*** Learner's posterior distributions. Plots show estimated amplitude across the scalp (32 electrodes). HDI: highest density intervals

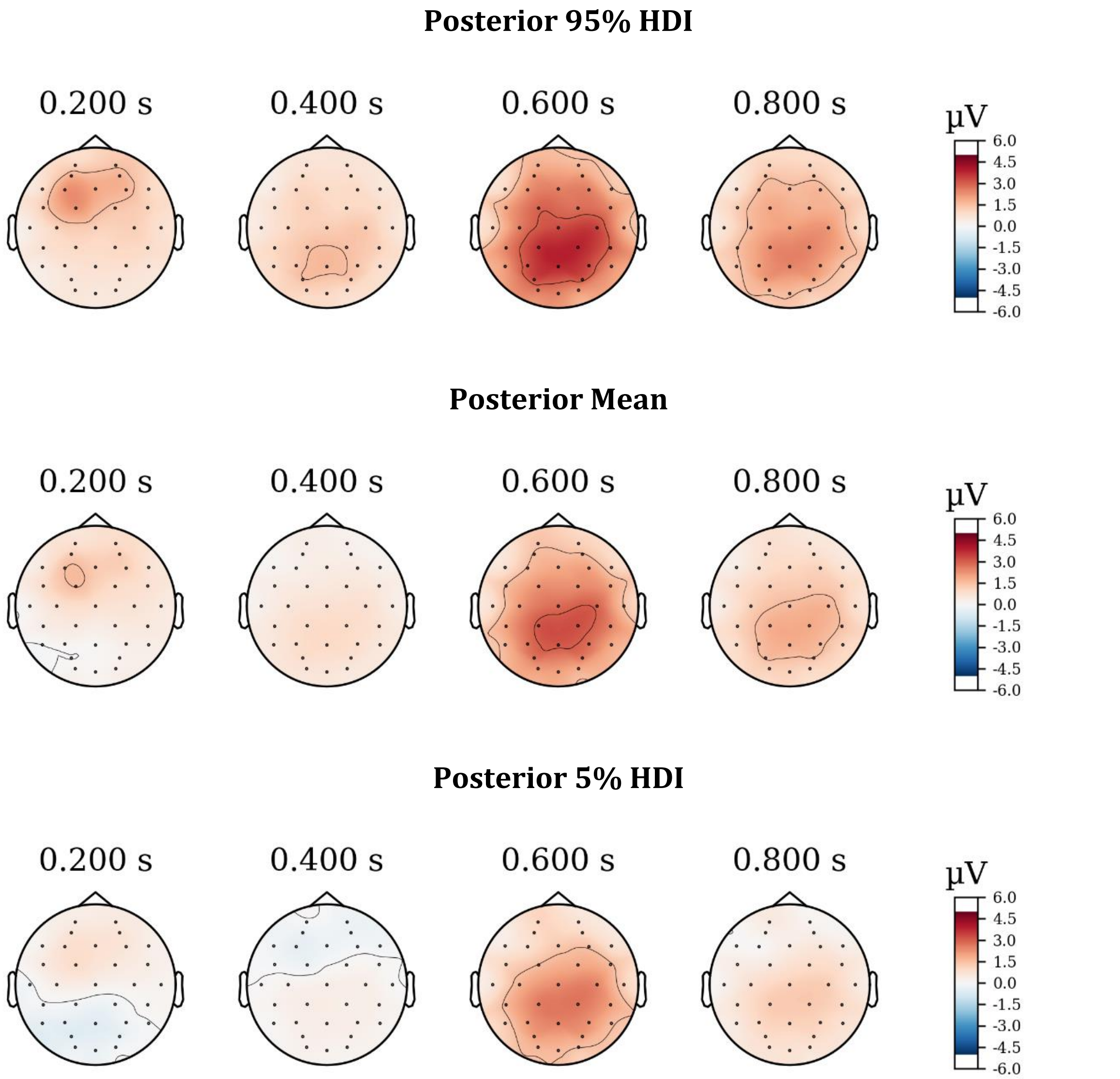


***Figure 4.*** Non-learner's posterior distributions. Plots show estimated amplitude across the scalp (32 electrodes). HDI: highest density intervals

*Model 3 Posterior Topomaps*

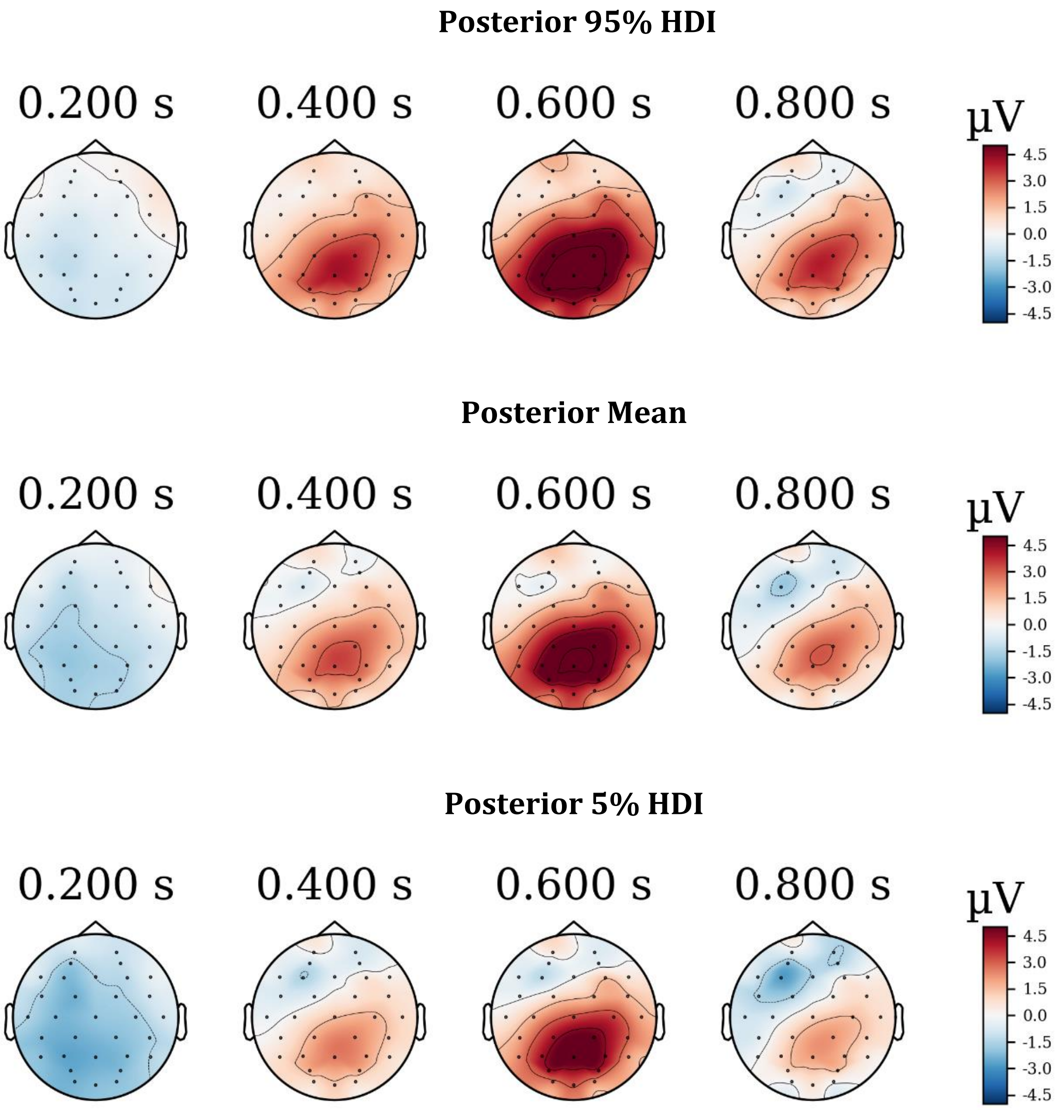


***Figure 5.*** Learner’s posterior distributions. Plots show estimated amplitude across the scalp (32 electrodes). HDI: highest density intervals

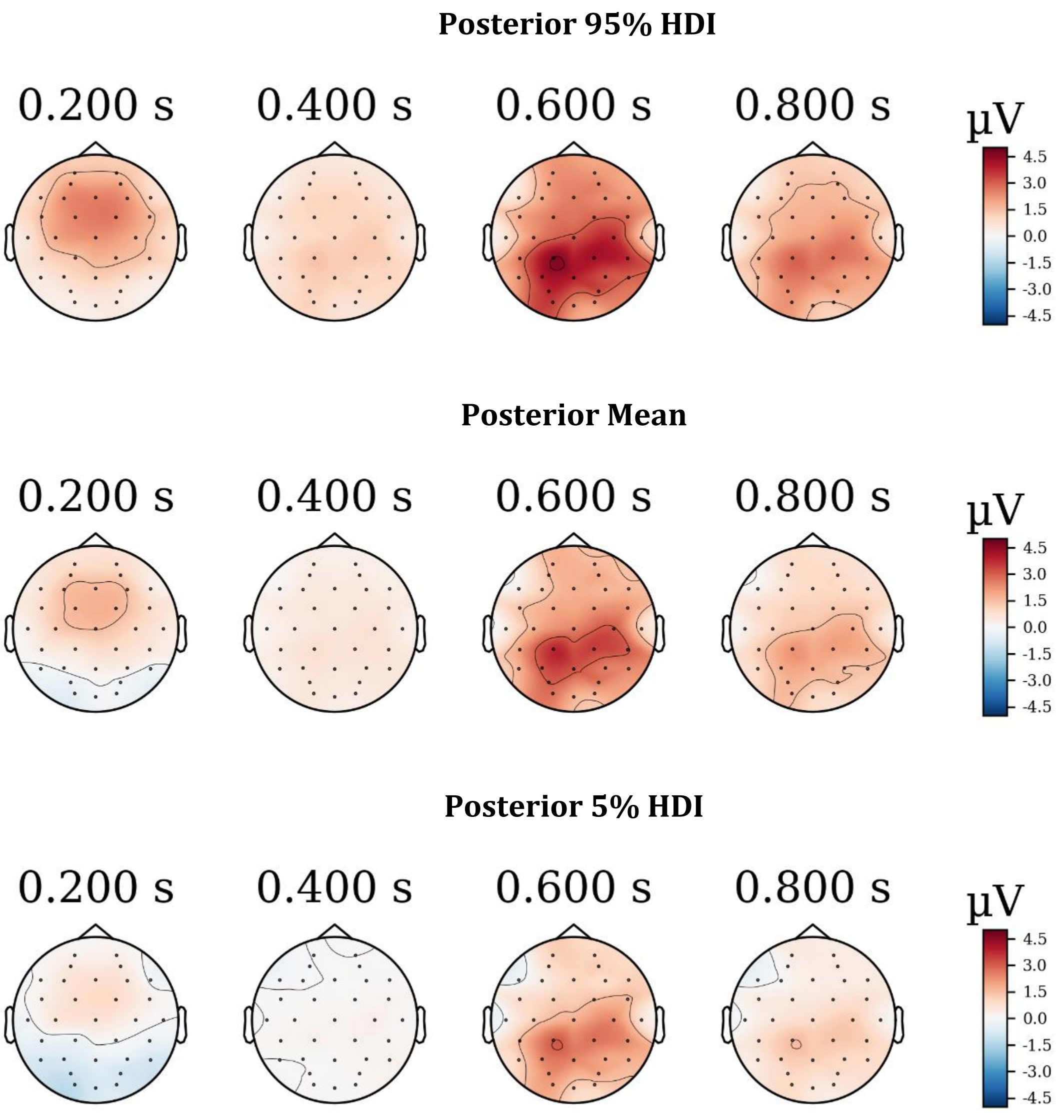


***Figure 6*.** Non-earner's posterior distributions. Plots show estimated amplitude across the scalp (32 electrodes). HDI: highest density intervals
