## Supplementary material for "Bayesian models for event-related potentials time-series and electrode correlations estimation": Annex 2

Present supplementary materials provide additional results, showcasing posterior predictive distributions.

### *Model 1 Posterior Predictions*

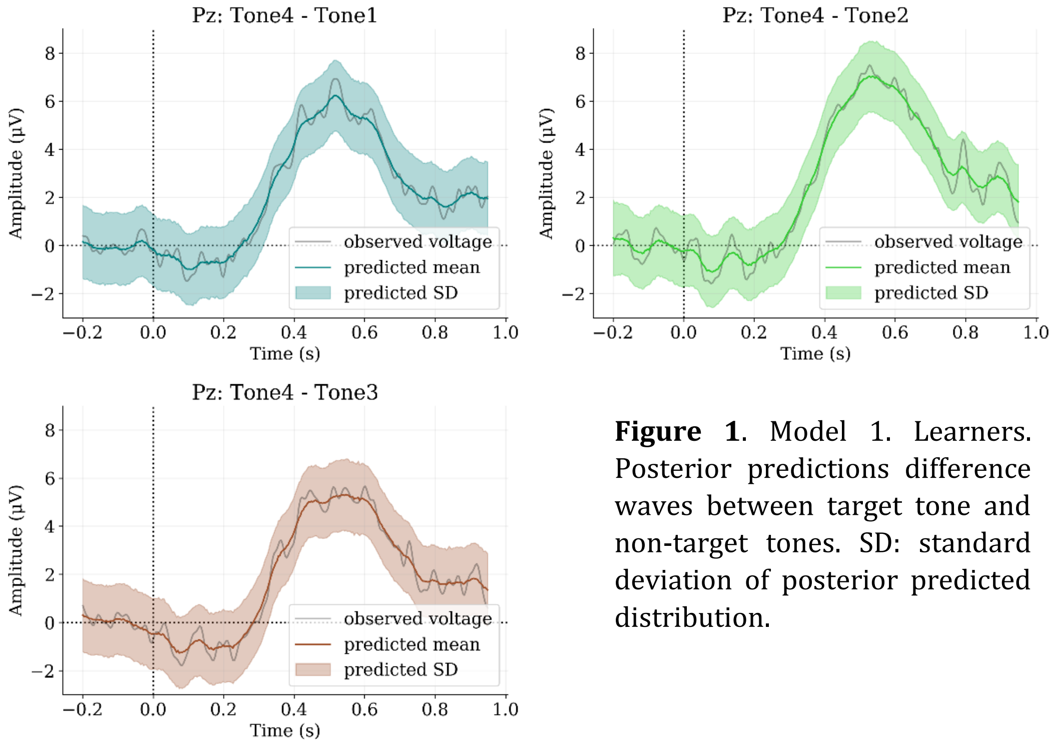


**Figure 1**. Model 1. Learners. Posterior predictions difference waves between target tone and non-target tones. SD: standard deviation of posterior predicted distribution.

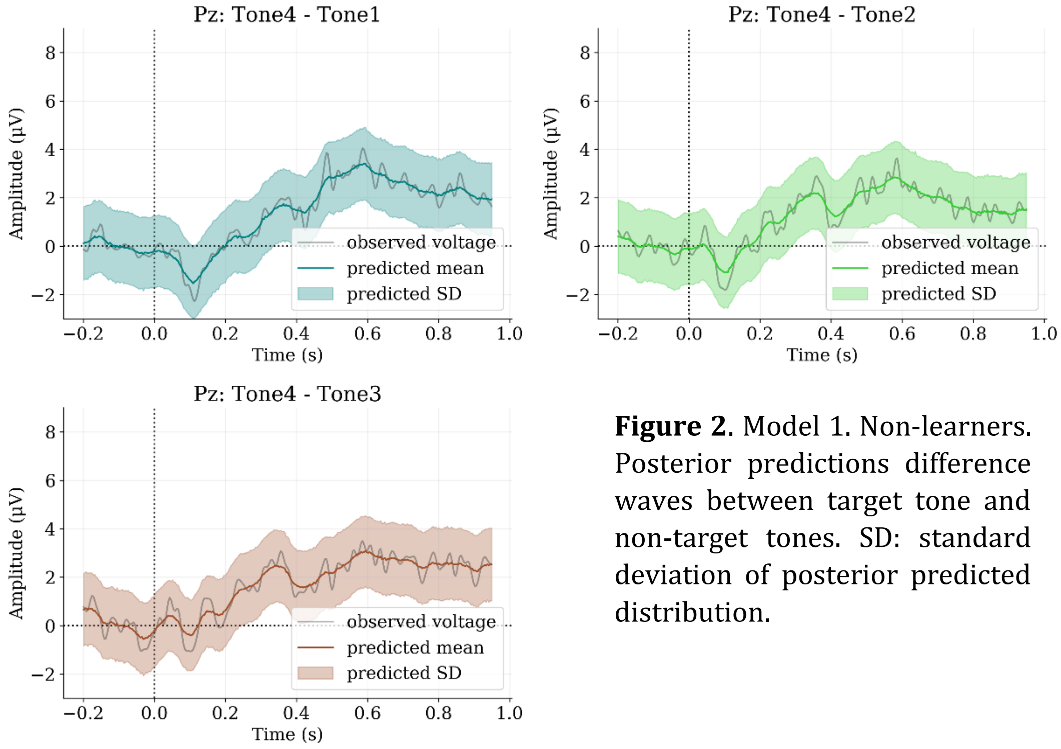


**Figure 2**. Model 1. Non-learners. Posterior predictions difference waves between target tone and non-target tones. SD: standard deviation of posterior predicted distribution.

*Model 2 Posterior Predictions*

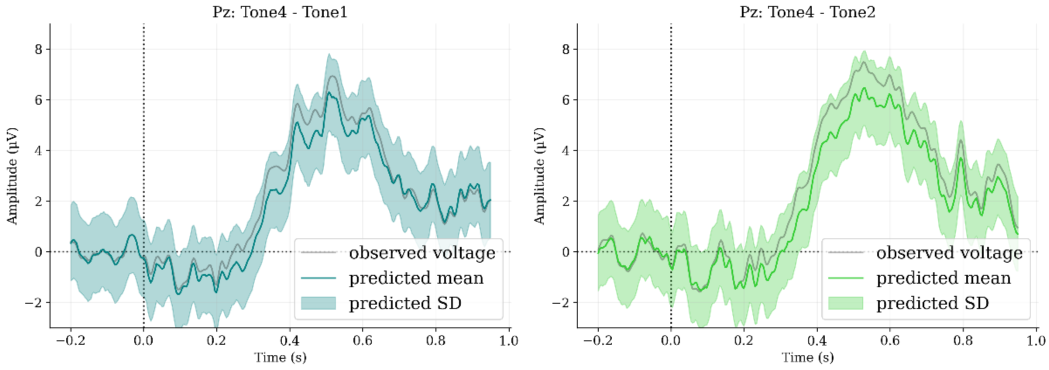


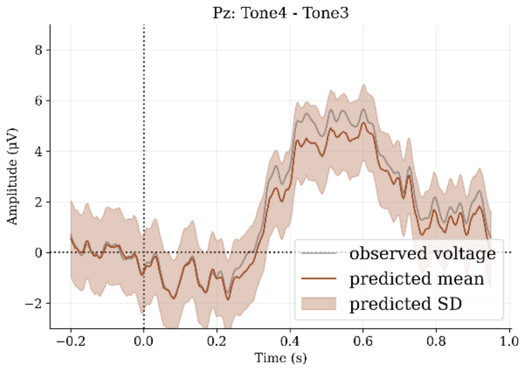


**Figure 3**. Model 2. Learners. Posterior predictions difference waves between target tone and non-target tones. SD: standard deviation of posterior predicted distribution.

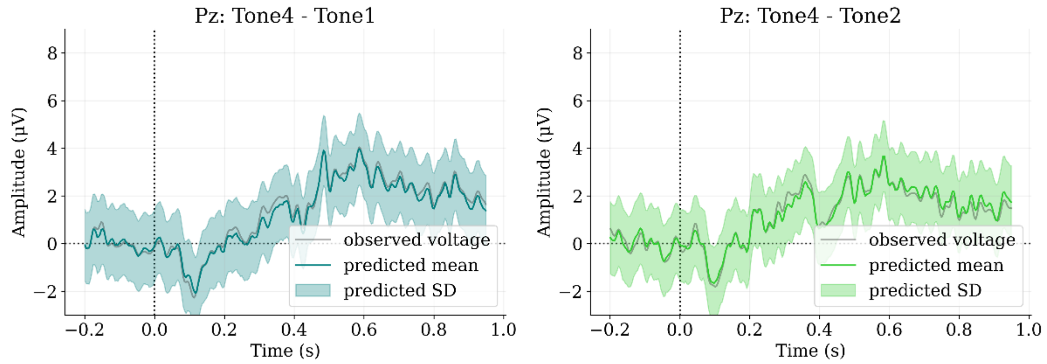


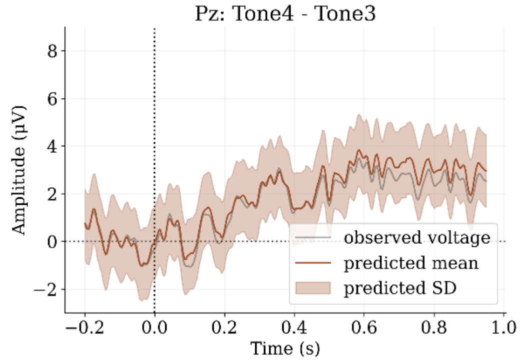


**Figure 4**. Model 2. Non-learners. Posterior predictions difference waves between target tone and non-target tones. SD: standard deviation of posterior predicted distribution.

*Model 3 Posterior Predictions*

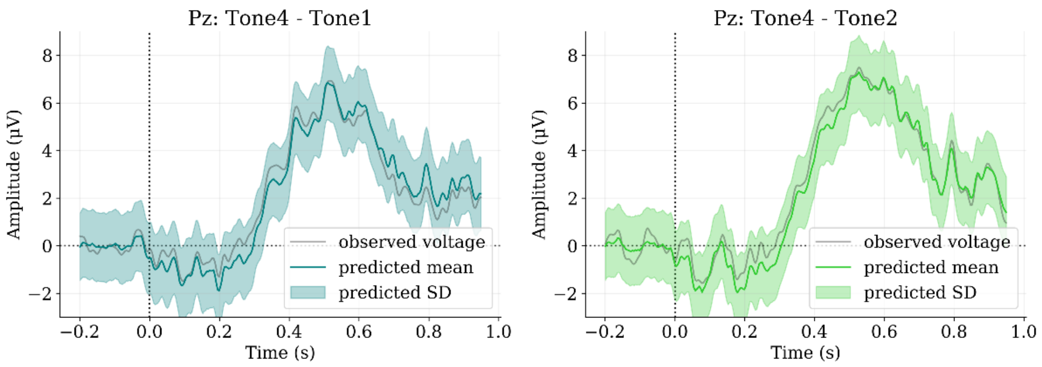


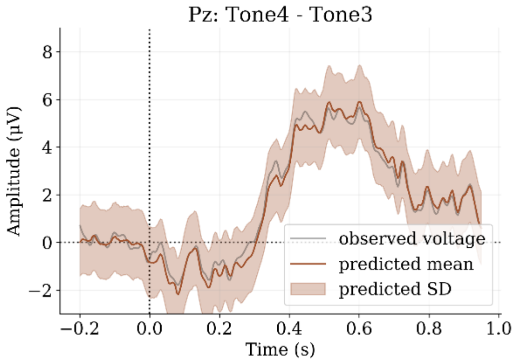


**Figure 5**. Model 3. Learners. Posterior predictions difference waves between target tone and non-target tones. SD: standard deviation of posterior predicted distribution.

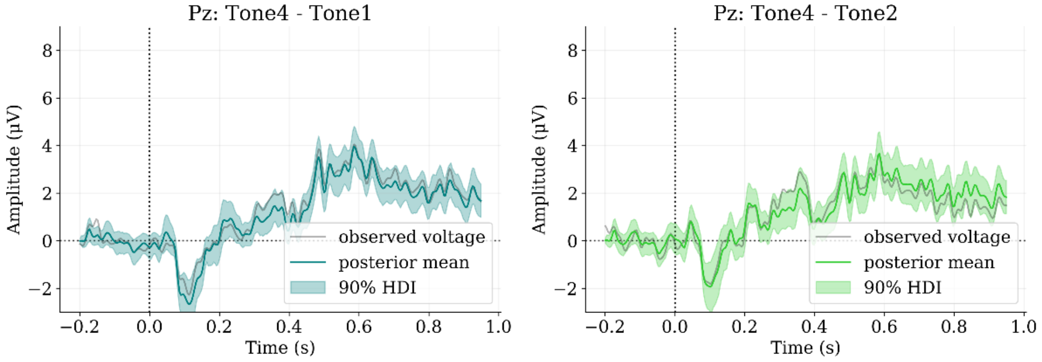


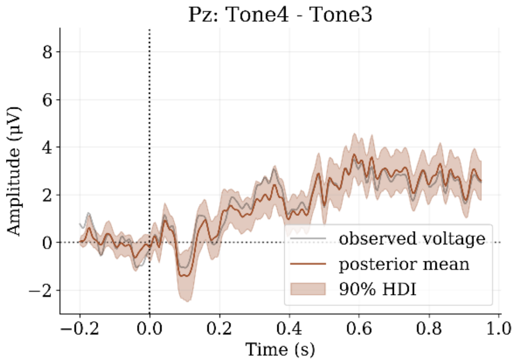


**Figure 6**. Model 3. Non-learners. Posterior predictions difference waves between target tone and non-target tones. SD: standard deviation of posterior predicted distribution.
